## Supplementary material for "Pseudo-pac site sequences used by phage P22 in generalized transduction of *Salmonella*": SupplFig1

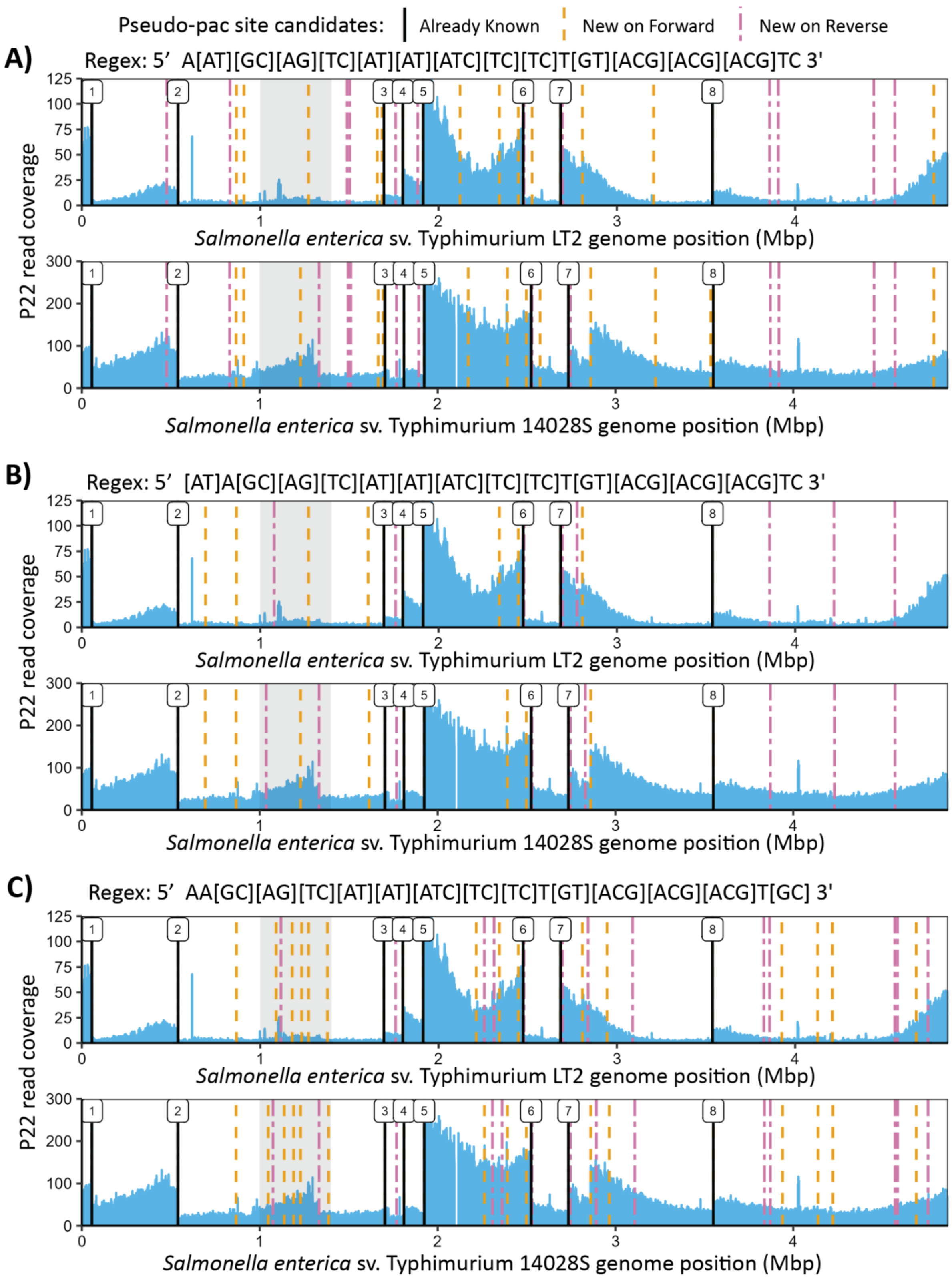


**Figure S1: Examples of *Salmonella* genome matches to other regular expression patterns**

A-C) Coverage plots of the *Salmonella enterica* sv. Typhimurium LT2 (LT2) and *Salmonella enterica* sv. Typhimurium 14028S (14028S) genomes with sequencing reads from purified P22. The additional generalized transduction site present in 14028S but not LT2 is shaded in grey. The regular expression (Regex) patterns used to search the *Salmonella* genomes for additional pseudo-pac sites are displayed above their associated plots. Black vertical lines indicate the locations of the pseudo-pac site sequences that were previously identified in this study. Orange and pink dashed lines indicate the locations of regular expression matches on the forward and reverse *Salmonella* genome strands, respectively.
