## Supplementary material for "Pseudo-pac site sequences used by phage P22 in generalized transduction of *Salmonella*": SupplText

### **Supplementary files:**

**Text:**

R code for regular expression pattern matching to genome sequences:

#Load packages

library(seqinr)

library(dplyr)

library(tidyr)

library(stringr)

#Import genome sequence fasta files

STLT2_seq <- read.fasta(file = "StyphimuriumLT2_genome.fasta")

ST14028S_seq <- read.fasta("Styphimurium14028S_genome.fasta")

\# Function to search genome sequences for custom regular expression matches

regex_function <- function(GenomeStringForward, RegexPattern, RegexPatternReverse){

regex_matches_forward <- c()

regex_matches_reverse <- c()

reverse_seq <- chartr("atgc", "tacg", GenomeStringForward)

X <- 1

Y <- 17

repeat{

site_subset <- GenomeStringForward[c(X:Y)]

site_string <- site_subset %>% toString()

site_string <- gsub(", ", "", site_string)

site_subset_comp <- reverse_seq[c(X:Y)]

site_string_comp <- site_subset_comp %>% toString()

site_string_comp <- gsub(", ", "", site_string_comp)

if (str_detect(site_string, RegexPattern) == TRUE) {

regex_matches_forward <- c(regex_matches_forward, X)

}

if (str_detect(site_string_comp, RegexPatternReverse) == TRUE) {

regex_matches_reverse <- c(regex_matches_reverse, X)

}

X <- X+1

Y <- Y+1

if (Y == length(GenomeStringForward)) break

}

regex_forward_matches <- data.frame(intercepts=regex_matches_forward, names=rep('New on Forward',length(regex_matches_forward)))

regex_reverse_matches <- data.frame(intercepts=regex_matches_reverse, names=rep('New on Reverse', length(regex_matches_reverse)))

return(list(regex_forward_matches, regex_reverse_matches))

}

#search LT2 genome sequence for 17bp sequence with same conservation as pseudo pac-site candidate sequences using the following regular expression (regex) pattern: aag[ag][tc][at][at][atc][tc][tc]t[gt][ac][ac][gca]tc

LT2_regex1 <- regex_function(STLT2_seq$NC_003197.2, "aag[ag][tc][at][at][atc][tc][tc]t[gt][acg][acg][gca]tc", "ct[acg][acg][acg][gt]t[tc][tc][atc][at][at][tc][ag]gaa")

new_on_forward_regex1LT2 <- LT2_regex1[[1]] #Positions of matches on forward genome strand

new_on_reverse_regex1LT2 <- LT2_regex1[[2]] #Positions of matches on reverse genome strand

#Search 14028S genome sequence for 17bp sequence with same conservation as pseudo pac-site candidate sequences using the following regular expression (regex) pattern: aag[ag][tc][at][at][atc][tc][tc]t[gt][ac][ac][gca]tc

ST_14028S_regex1 <- regex_function(ST14028S_seq$NC_016856.1, "aag[ag][tc][at][at][atc][tc][tc]t[gt][acg][acg][gca]tc", "ct[acg][acg][acg][gt]t[tc][tc][atc][at][at][tc][ag]gaa")

new_on_forward_regex114028S <- ST_14028S_regex1[[1]] #Positions of matches on forward genome strand

new_on_reverse_regex114028S <- ST_14028S_regex1[[2]] #Positions of matches on reverse genome strand

#Repeat the regex_function() call on both the LT2 and 14028S genomes for other regex patterns:

#aa[cg][ag][tc][at][at][atc][tc][tc]t[gt][acg][acg][gca]tc

#[at]a[cg][ag][tc][at][at][atc][tc][tc]t[gt][acg][acg][gca]tc

#a[at][cg][ag][tc][at][at][atc][tc][tc]t[gt][acg][acg][gca]tc
